# The RNA binding protein Nab2 regulates the proteome of the developing *Drosophila* brain

**DOI:** 10.1101/2020.12.10.419846

**Authors:** Edwin B. Corgiat, Sara M. List, J. Christopher Rounds, Anita H. Corbett, Kenneth H. Moberg

## Abstract

The human *ZC3H14* gene, which encodes a ubiquitously expressed polyadenosine zinc finger RNA binding protein, is mutated in an inherited form of autosomal recessive, non-syndromic intellectual disability. To gain insight into ZC3H14 neurological functions, we previously developed a *Drosophila melanogaster* model of ZC3H14 loss by deleting the fly ortholog, Nab2. Studies in this invertebrate model reveal that Nab2 controls final patterns of neuron projection within fully developed adult brains. Here, we examine earlier pupal stages and define roles for Nab2 in controlling the dynamic growth of axons into the developing brain mushroom bodies (MBs), which support olfactory learning and memory, and in regulating abundance of a small fraction of the total brain proteome, a portion of which is rescued by overexpression of *Nab2* specifically in brain neurons. The group of Nab2-regulated brain proteins, identified by quantitative proteomic analysis, includes the microtubule binding protein Futsch, the neuronal Ig-family transmembrane protein Turtle, the glial:neuron adhesion protein Contactin, the RacGAP Tumbleweed, and the planar cell polarity factor Van Gogh, which collectively link Nab2 to a the processes of brain morphogenesis, neuroblast proliferation, circadian sleep/wake cycles, and synaptic development. Overall, these data indicate that Nab2 controls abundance of a subset of brain proteins during the active process of wiring the pupal brain mushroom body, and thus provide a window into potentially conserved functions of the Nab2/ZC3H14 RNA binding proteins in neurodevelopment and function.

## Introduction

Neurons develop complex architectures that allow them to function within massive interconnected networks that transmit electrochemical signals among thousands of other neurons in a shared circuit. The polarized morphology of neurons is particularly unique, with each cell containing axon and dendrite projections that can extend over enormous distances relative to the size of the cell body. Axonal growth and guidance is largely directed through the growth cone, which responds to guidance cues to steer the axon (1, 2). This axonal guidance is regulated in part by local translation of mRNAs within the growth cone that modifies the local proteome. This process of local translation, which relies on pre-delivery of mRNAs to the axon tip, facilitates rapid shifts in translation in response to extracellular cues that would otherwise be limited by distance from the nucleus and relatively slow speed of intracellular transport (1–4). The local translation of mRNAs in distal neuronal projections is critical for proper development of the nervous system (1, 3) but poses many biological challenges, including the need to maintain mRNAs in a translationally repressed state during transport from the nuclear periphery to distal sites where regulated translation must occur (2, 4). RNA binding proteins (RBPs) play a major role in this process (4).

RBPs play critical roles in regulating temporal and spatial expression of numerous mRNAs that encode proteins with roles in neuronal function (5). Although RBPs play broadly important roles in regulating multiple steps in gene expression that are shared by all cell types, mutations in genes encoding RBPs often result in tissue- or cell-type specific diseases (2, 4, 6–10). A large number of these RBP-linked diseases include significant neurologic impairments, which likely reflects an enhanced reliance on post-transcriptional mechanisms to pattern spatiotemporal gene expression over the long distances that neurons extend (1, 11, 12). This dependence on RBP-based mechanisms of gene expression is exemplified by disease-causing mutations in the genes (4) encoding the fragile X mental retardation protein (FMRP) (13), survival of motor neuron protein (SMN) (14), and TAR DNA binding protein 43 (TDP-43) (11). Mutations in the *ZC3H14* gene, which encodes a zinc finger RBP (zinc finger CysCysCysHis [CCCH]-type 14), cause neurological defects that broadly resemble those associated with these more extensively characterized RBPs (4, 15).

The human *ZC3H14* gene encodes a ubiquitously expressed polyadenosine RNA binding protein that is lost in a heritable non-syndromic form of intellectual disability (15). The *Drosophila* ZC3H14 homolog, Nab2, has provided an excellent model to probe the function of ZC3H14/Nab2 in neurons (16–18). *Nab2* deletion in flies results in defects in locomotion and neuromorphology that are rescued by neuronal specific re-expression of Nab2 (17). Neuronal specific expression of human ZC3H14 partially rescues many of the Nab2 null phenotypes, demonstrating a high level of functional conservation between ZC3H14 and Nab2 (17–19).

Nab2 and its orthologs are found primarily in the nucleus at steady-state (20–25), but evidence shows that these proteins can shuttle between the nucleus and cytoplasm (22, 23, 26). A small pool of Nab2 is detected in the axonal and dendritic cytoplasm of neurons (15–17, 20, 21) raising the possibility that Nab2 has both nuclear and cytoplasmic roles in neurons. Multiple studies in a variety of model organisms have defined key roles for Nab2 in pre-mRNA processing events within the nucleus, including regulation of splicing events, transcript termination, and control of poly(A) tail length (19, 27–30). Additional studies localize Nab2 within cytoplasmic mRNA ribonucleoprotein particles and imply roles in translational repression, likely mediated in part through interactions with FMRP (20–23, 28). Ultimately, all of these post-transcriptional regulatory events are likely to alter levels of key proteins that are critical for proper neuronal function.

At a morphological level, zygotic deficiency for Nab2 produces structural defects in the adult brain mushroom bodies (MBs) (17), twin neuropil structures that mirror across the brain midline and are required for olfactory learning and memory (17, 31, 32). The MBs are formed of five lobes: γ, α, α’, β, and β’ (**Figure 1A**) (33, 34). In the fully formed adult brain, *Nab2* null neurons fail to project axons into the α-lobe and β-lobe axons inappropriately cross the midline into the contralateral hemisphere (17, 20). These findings implicate Nab2 in developmental control of axonogenesis and growth cone guidance. MB development begins in the larval stage with neuroblast pools that project axons into nascent γ-lobes (33–37). During the subsequent pupal stage, these γ-lobes are pruned back, and α and β-axons begin to project into their corresponding tracks (33–37). By 24 hours after pupal formation (APF), α and β-lobes have formed their initial structure and are being thickened by new axons that project through the core of the bundle. This process continues through ~72 hours APF, when the α and β-lobes are fully formed (33, 35). The effect of *Nab2* alleles on final α and β-lobe structure in the adult brain implies a role for the Nab2 RBP in axon projection and guidance during early pupal stages(17, 20, 34–36).

**Figure 1.**
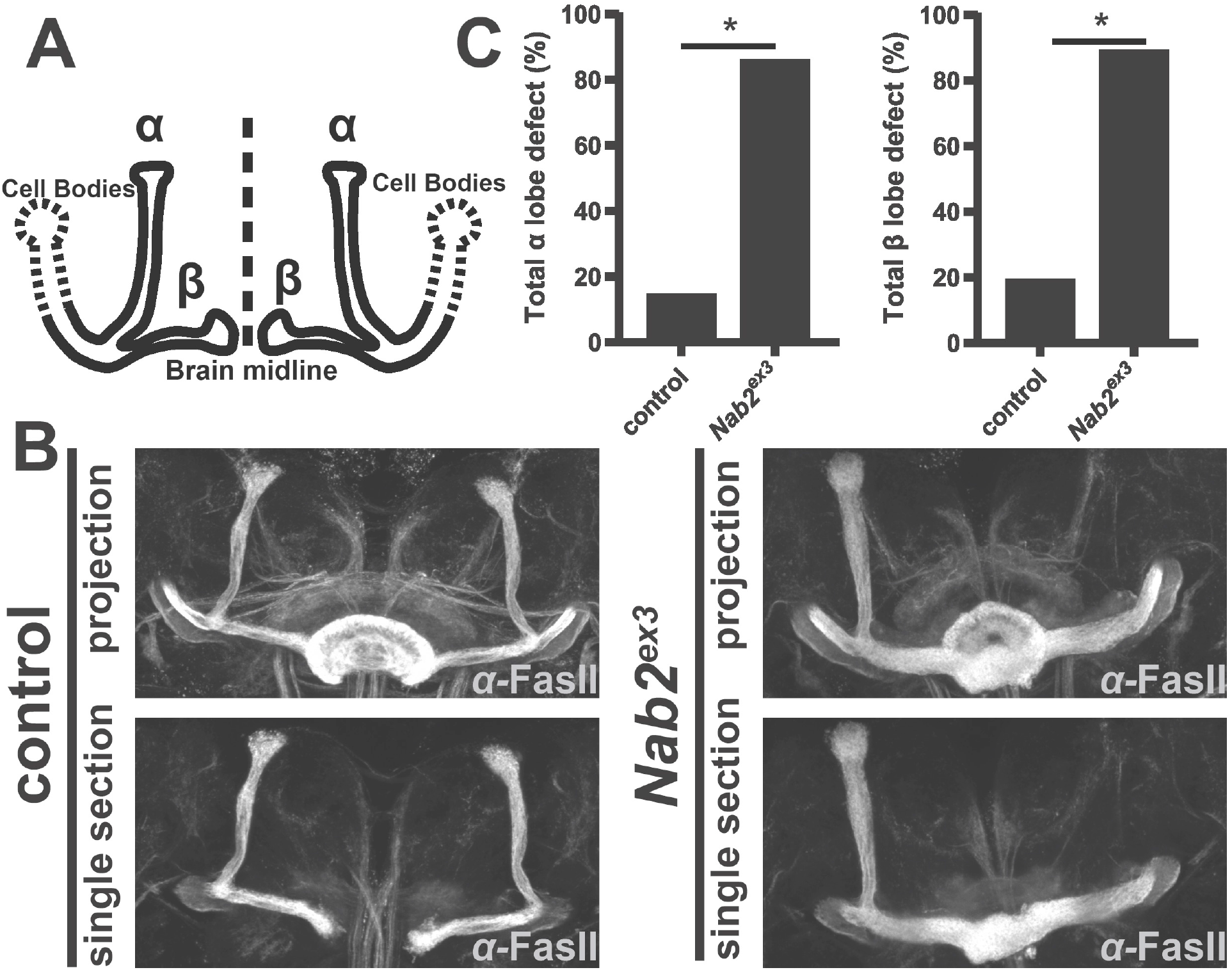
Nab2 is required during pupal development for proper neuro-morphological patterning of the mushroom bodies. **(A)** Diagram of the *Drosophila* mushroom body depicting cell bodies (dashed lines) projecting axons that bundle to make the dorsal (α) and medial (β) lobes that are mirrored across the brain midline (dashed line). **(B)** Fasciclin II (FasII) antibody staining of *control (C155>Gal4. w^1118^*) and *Nab2^ex3^ (C155>Gal4;;Nab2^ex3^*) 48-72hr after pupal formation brains. Confocal images show maximum intensity Z-stack projections (projection) which display full mushroom bodies and single transverse plane sections (single section) which display midline crossing of β-lobe axons. Imaging reveals that *control* rarely shows defects in α and β-lobes while *Nab2^ex3^* brains often have thinning or loss of the α-lobes and β-lobes that project across the midline into the contralateral hemisphere resulting in fusion of the lobes or occasionally loss of β-lobes. The ellipsoid body (donut shaped structure at the brain midline) is visible in maximum intensity projection images which masks the β-lobe status, so single section images are included for clarity. **(C)** Quantification of the frequency of *control* and *Nab2^ex3^* (left) total α-lobe defect (thinning or missing α-lobe) or (right) total β-lobe defect (fusion or missing β-lobes) using the scoring system as described in Experimental Procedures. * indicates p<0.05.

Here, we exploit the predictable time course of brain development to perform a temporally coupled analysis of Nab2 regulation of the pupal brain proteome by quantitative proteomic analysis and pair this with a parallel analysis of axon projection into the forming pupal MBs. We find that Nab2 loss disrupts α and β-axon projection in the pupal MBs coincident with significant increases in the steady-state abundance of proteins that are highly enriched for roles in neurodevelopment, neuronal and glial metabolism, axon guidance, and trans-synaptic signaling. Complementary analysis of neuronal specific Nab2-overexpressing brains confirms that a subset of these proteins also change abundance in response to excess Nab2. In sum, this paired morphological-proteomic analysis provides strong evidence that Nab2 is required to control the abundance of proteins with critical roles in *Drosophila* neurons that may play conserved roles in humans.

## Results

### Nab2 loss disrupts axon projection into the forming pupal mushroom bodies

Our prior finding that loss of Nab2 impairs mushroom body (MB) neuromorphology in the mature adult *Drosophila* brain (4, 6, 7) suggests a role for Nab2 in MB morphogenesis in the preceding pupal phase. Consistent with this idea, serial optical sectioning of α-FasII-stained *Nab2^ex3^* (i.e. zygotic null) and *control* brains 48-72 hours after pupal formation (APF) reveals thinning or missing α-lobes and β-lobes that project and fuse across the midline that are not present to the same extent in *control* brains (**Figure 1A-B**). The 48-72 hr APF time window coincides with a midpoint in projection and guidance of α and β-lobes. At this stage, *control* brains show incompletely formed α and β-lobes with a low degree of defects (13% and 18%, respectively) while *Nab2^ex3^* brains already display a high rate of missing/thinning α-lobes and fused/missing β-lobes (both 85%) (**Figure 1C**). These data indicate that *Nab2* is required during pupal projection and guidance of the mushroom body axons, raising the question of how loss of the Nab2 RBP affects the pupal brain proteome.

### Quantitative proteomic analysis of developmentally timed pupal brains

Nab2 has been identified as a component of cytoplasmic ribonucleoprotein particles (RNPs) linked to mRNA trafficking and translation (20, 23, 25), and as a nuclear component of post-transcriptional complexes (20, 21) that control mRNA splicing (22, 27, 28), transcription termination (30), and polyadenylation (16). To explore how *Drosophila* Nab2 affects the mRNA-derived proteome in the developing pupal brain, global label-free LC-MS/MS was performed on dissected 24hr APF brains of *control (C155>Gal4, w^1118^*), mutant *Nab2^ex3^ (C155>Gal4;;Nab2^ex3^*), and neuronal specific *Nab2* overexpression (*Nab2 oe*) (*C155>Gal4;Nab2^EP3716^;Nab2^ex3^*) animals as illustrated in **Figure 2**. The final genotype takes advantage of our prior work showing that reexpression of *Nab2* solely in the neurons of otherwise *Nab2^ex3^* mutant animals rescues many *Nab2^ex3^* organismal phenotypes (15, 17) and provides an opportunity to identify neuronal proteins that are also rescued in this *Nab2^ex3^ oe* paradigm. We employed 24h APF brains for proteomic analysis in order to capture the developmental window during which MB defects resulting from Nab2 loss form.

**Figure 2.**
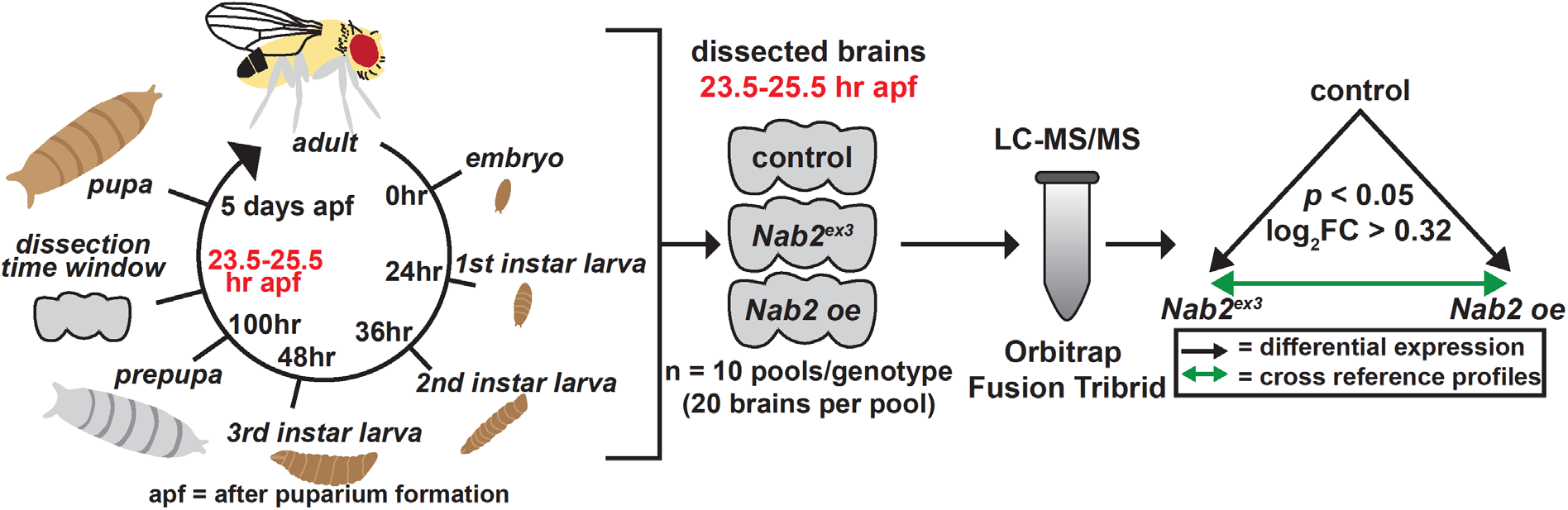
Study design and analytic approach for quantitative proteomic analysis of *Drosophila* pupal brains. A workflow summary showing dissection window, experimental design, and analysis. The *Drosophila* life cycle with developmental stage and hours of development depicted with the dissection time window (23.25-25.5 hr APF) in **red, left**. There were 600 developmentally timed brain samples which were pooled by genotype: *control* (*C155>Gal4, w^1118^*); Nab2 zygotic null (*Nab2^ex3^* = *C155>Gal4;;Nab2^ex3^); Nab2 overexpression in neurons (Nab2 oe* = *C155>Gal4;Nab2^EP3716^;Nab2^ex3^*) and by sex resulting in 30 individual pools, **center**. Each sample pool was processed, analyzed using an Orbitrap Fusion Tribrid Mass Spectrometer, and quantified using MaxQuant against the *D. melanogaster* Uniprot database, **center**. Arrows depict the performed analyses. Differential protein abundance of *Nab2^ex3^* and *Nab2 oe* brains was calculated with an FDR adjusted p-value (**black** arrows), and then second-degree analyses cross referencing the *Nab2^ex3^* and *Nab2 oe* proteomic profiles (**green** arrows)**, right**.

Unbiased principal component analysis (PCA) of the proteomic data reveals three distinct clusters (**Figure 3A**). The thirty samples cluster by genotype, forming three clusters with ten samples in each cluster. Mass spectrometry was carried out for ten biological replicates per genotype, with five male samples and five female samples analyzed separately. Subsequent simple linear regression modelling of the data obtained indicated that male and female samples could be combined for analyses adding power. These combined datasets (n=10 per genotype) were used for subsequent analyses.

**Figure 3.**
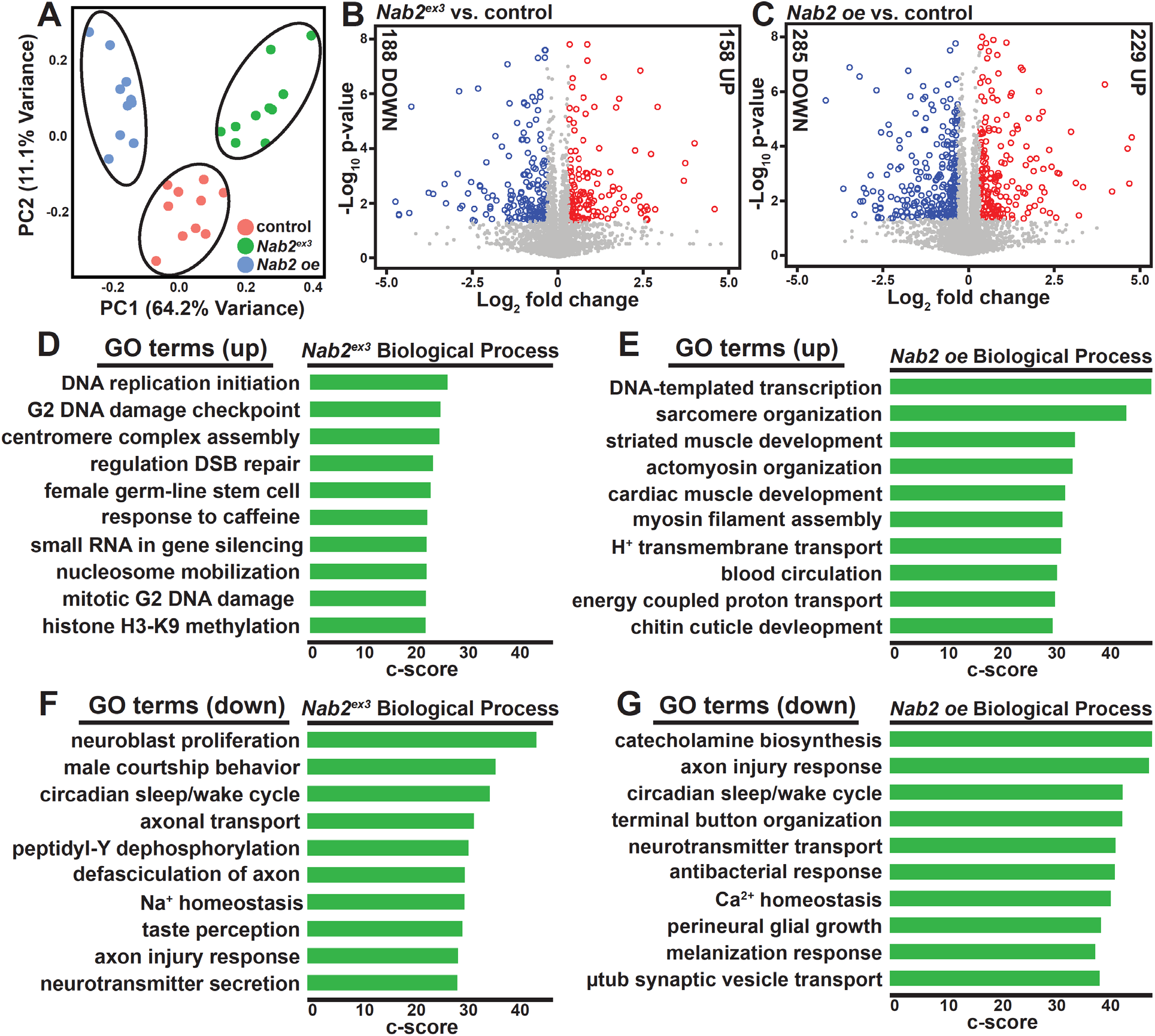
Quantitative proteomic analysis of developmentally timed pupal brains reveals a role for Nab2 in neurodevelopment. **(A)** Principal component analysis (PCA) of proteomic data from 24hr after pupal formation *Drosophila* brains from ten biological replicates of *control, Nab2^ex3^*, and *Nab2 oe* flies (*control* = *C155>Gal4, w^1118^; Nab2^ex3^* = *C155>Gal4;;Nab2^ex3^; Nab2 oe* = *C155>Gal4;Nab2^EP3716^; Nab2^ex3^*) show results cluster based on genotype and that *Nab2^ex3^* and *Nab2 oe* are distinct from *control* and each other. **(B,C)** Volcano plots show proteins differentially expressed in each *Nab2* genotype [(B) *Nab2^ex3^* (346; 188 down and 158 up) and **(C)** *Nab2 oe* (514; 285 down and 229 up)] compared to *control* (adj p-val<0.05). Protein abundance change (Down or Up) indicated on each side of the plot (log_2_ average protein abundance: **grey** = n.s., **blue** ≤ −0.32, **red** ≥ 0.32). **(D-G)** The enriched terms from FlyEnrichr database for Biological Process are shown for proteins increased log_2_ fold-change <0.32 in **(D)** *Nab2^ex3^* and (E) *Nab2 oe* and decreased log_2_ fold-change >-0.32 in **(F)** *Nab2^ex3^* and **(G)** *Nab2 oe*. The bars shown correspond to the top ten c-scores (= ln(adj p-val) * z-score) in each dataset (adj. p-val<0.05).

As previous studies suggest Nab2 can function as a translational repressor (20, 21), the most direct Nab2 targets could be expected to increase in abundance. However, factors that decrease in protein abundance may also be phenotypically significant in the *Nab2^ex3^* genotype. Of the 4302 proteins identified as ‘present’ across all three groups, 346 proteins (approximately 8% of total proteins detected) are differentially expressed in the *Nab2^ex3^* brains vs *control* brains (**Figure 3B**) (full dataset available at ProteomeXchange Consortium via PRIDE under the accession #PXD022984). Within this group, 158 proteins score ≥0.8 log_2_ fold change increase in expression (five most elevated: CG1910, Got1, Ida, Mtp, and Wwox) and 188 proteins score ≤-0.8 log_2_ fold change decrease in expression, with Nab2 among the top five most decreased (remaining four: Pglym78, Mkk4, Cortactin, and Psa) (**Figure 3B**). Of the 514 proteins differentially expressed in *Nab2 oe* brains relative to *control* brains (approximately 12% of total proteins detected), 229 proteins score ≥0.8 log_2_ fold change increase (five most elevated: CG1910, Ccp84Ae, Ida, Ccp84Ag, and Alien) and 285 proteins scored ≤-0.8 log_2_ fold change decrease (five most decreased: Pglym, Mkk4, Cortactin, Gnmt, and CG34280) (**Figure 3C**). Comparing the differentially expressed proteins from *Nab2^ex3^* brains to a previously reported proteomic dataset generated from hippocampi of P0 *Zc3h14* knockout (*Zc3h14^Δex13/Δex13^*) mice (21), reveals six proteomic changes shared between flies and mice (**Table 1**). These conserved changes may give insight into conserved effects of Nab2/ZC3H14 loss across flies and mice.

**Table 1.** Evolutionarily conserved proteomic changes between *Nab2^ex3^* flies and *Zc3h14^Δex13/Δex13^* mice.

| Fly Protein Symbol | $\log_2(Nab2^{ex3}/Cont)$ | Mouse Protein Symbol |
| --- | --- | --- |
| Wwox | 7.5 | Wwox |
| Asrij | 0.6 | Ociad1 |
| Sec71 | -1.3 | Psd3 |
| Vang | -2.3 | Vangl2 |
| CG6693 | -2.8 | Dnajc9 |
| X11Lbeta | -4.7 | Apba1 |

A FlyEnrichr analysis (FlyEnrichr: amp.pharm.mssm.edu/FlyEnrichr/) for process enrichment within the significantly changed *Nab2^ex3^* and *Nab2 oe* proteomic profiles reveals that proteins increased in *Nab2^ex3^* represent biological processes involved in genome maintenance (e.g. DNA replication initiation, G2 DNA damage checkpoint, centromere complex assembly) and development (e.g. female germline stem cell) (**Figure 3D**), while proteins increased in *Nab2 oe* represent processes related to development (e.g. striated muscle development, cuticle development) and muscle organization (e.g. sarcomere organization, myosin filament assembly) (**Figure 3E**). The profile of decreased proteins in *Nab2^ex3^* and *Nab2 oe* proteomic profiles are both strongly enriched for processes linked to neurodevelopment, synaptic function, and brain maintenance (**Figure 3F, G**). Within the *Nab2^ex3^* dataset, decreased proteins are enriched for the processes of neuroblast proliferation, circadian sleep/wake cycle, and axonal transport (**Figure 3F**). Within the *Nab2 oe* dataset, decreased proteins are enriched for the processes of axon injury response, circadian sleep/wake cycle, and neurotransmitter transport (**Figure 3G**).

Comparison of the *Nab2^ex3^* and *Nab2 oe* brain proteomes provides some significant insights. First, although there are significant differences in the identity (**Figure 4A**) and direction of effect (**Table 2**) on affected proteins, there are significant correlations between these two datasets (**Figure 4B**). A total of 195 proteins are changed in both *Nab2^ex3^* and *Nab2 oe* brains (**Figure 4A**) and are highly correlated with one another (R=0.86, p<2.2^-16^; **Figure 4B**). Of these 195 shared proteins, a large fraction (184 out of 195, approximately 94%) change abundance in the same direction (**Table 2**). Second, each dataset has proteins changes that are unique to the *Nab2* loss (*Nab2^ex3^*) or *Nab2* neuronal-overexpression (*Nab2 oe*) genotypes: 152 proteins are changed exclusively in *Nab2^ex3^* brains, and 311 proteins are changed exclusively in the *Nab2 oe* brains. As general overexpression of *Nab2* is more lethal than zygotic *Nab2* loss (15), the 311 changes unique to *Nab2 oe* may represent dominant effects of excess Nab2. However, the 152 proteins that are significantly changed only in *Nab2^ex3^* brains, and not in the *Nab2 oe* genotype (which is in the *Nab2^ex3^* background), are thus rescued by reexpression of wildtype Nab2 in *Nab2^ex3^* brain neurons. These differences in *Nab2^ex3^* and *Nab2 oe* proteomic profiles are also reflected in the FlyEnrichr analysis, which reveals 172 terms unique to *Nab2^ex3^* and 999 unique to *Nab2 oe* (**Figure 4C**).

**Figure 4.**
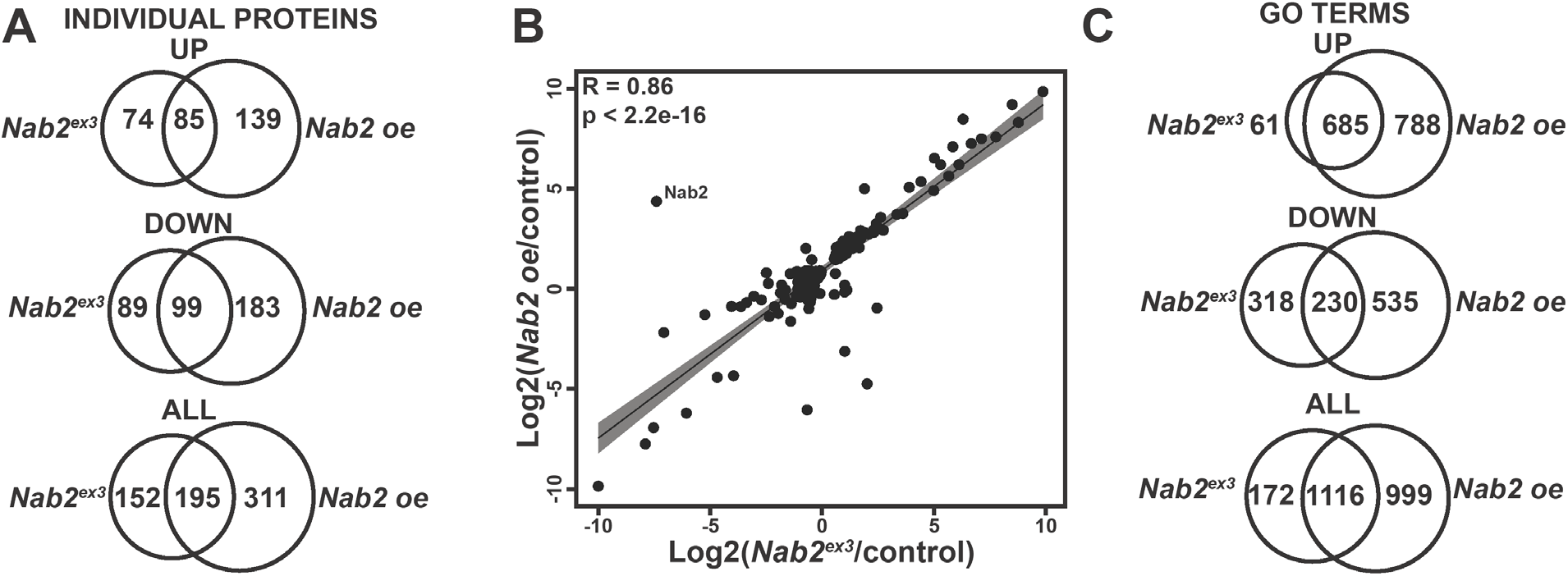
*Nab2^ex3^* and *Nab2 oe* brains display distinct proteomic profiles but have consistent changes among shared proteins. **(A)** Venn diagrams illustrating the number of individual proteins that show statistically significant changes, which are shared or unique to *Nab2^ex3^* and *Nab2 oe* that (top) increase or (middle) decrease in protein abundance (*Nab2^ex3^* relative to *control* and *Nab2 oe* relative to control) or (bottom) all abundance changes. **(B)** A correlation curve comparing the changes in protein abundance for proteins changed in both *Nab2^ex3^* and *Nab2 oe* relative to *control* was produced by plotting on a logarithmic scale. Results show that the shared changes (195) observed are highly correlated (R = 0.86, p < 2.2e-16, Pearson coefficient) in magnitude and direction. Regression line plotted in **black** with 95% confidence interval depicted by **grey** shading. Nab2 is expected to change in direction and magnitude between *Nab2^ex3^* and *Nab2 oe* and is annotated on the plot. **(C)** Venn diagrams illustrating the number of GO Biological Process terms enriched in *Nab2^ex3^* and *Nab2 oe* that are shared or unique based on the subset of proteins that (top) increase or (middle) decrease protein abundance or (bottom) all abundance changes.

**Table 2.** *Nab2^ex3^* and *Nab2 oe* shared proteins that change in different directions

| **Protein**<br>**Symbol** | **$\log_2(Nab2^{ex3}/Cont)$** | **$-\log_{10}(p\text{-value})$** | **$\log_2(Nab2oe/Cont)$** | **$-\log_{10}(p\text{-value})$** |
| --- | --- | --- | --- | --- |
| Nab2 | -8.4 | 8.3 | 3.9 | 6.2 |
| Hml | -1.1 | 2.0 | 1.0 | 3.3 |
| Mhc | -0.8 | 1.8 | 0.3 | 4.0 |
| LamC | 0.3 | 2.2 | -1.8 | 1.6 |
| CG15369 | 0.4 | 2.3 | -0.5 | 2.1 |
| Sgs7 | 0.8 | 1.9 | -1.7 | 2.2 |
| Sgs5 | 0.8 | 3.1 | -1.2 | 3.1 |
| Sgs3 | 0.8 | 1.3 | -5.3 | 2.7 |
| Sgs8 | 0.9 | 2.2 | -1.5 | 3.0 |
| Eig71Ed | 1.9 | 2.2 | -7.3 | 2.0 |
| sls | 2.4 | 6.8 | -2.6 | 1.3 |

The unique and shared changes in the *Nab2^ex3^* and *Nab2 oe* datasets were further divided in *increased* and *decreased* datasets, and then subjected to FlyEnrichr analysis. Protein increases unique to *Nab2^ex3^* represent processes involved in metabolism (**Figure 5A**), while *Nab2 oe* unique increases represent processes involved in tissue development and organization (**Figure 5B**). The increases common to *Nab2^ex3^* and *Nab2 oe* are enriched in processes involved in genome maintenance and development (**Figure 5C**). A chord plot of biological process GO terms relating to RNA processing and neurodevelopment highlights proteins *increased* in both *Nab2* and *Nab2 oe* datasets (**Figure 5D**). Among these are the glial:neuronal adhesion protein Contactin (Cont), the planar cell polarity (PCP) accessory protein A-kinase anchor protein 200 (Akap200), and the condensin subunit Gluon (Glu), and the regulator of neuroblast division Polo (**Figure 5D**).

**Figure 5.**
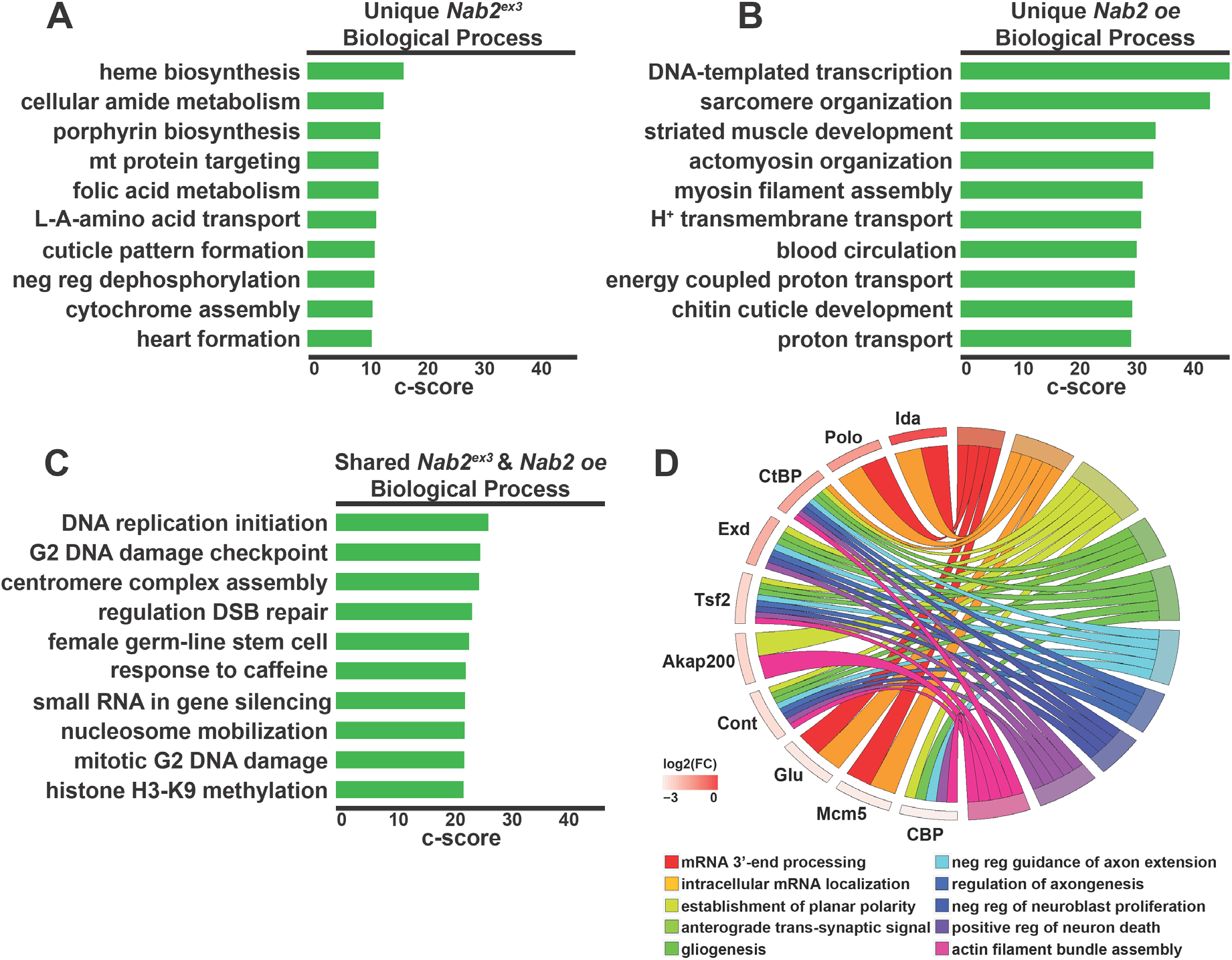
Proteins increased in abundance in *Nab2^ex3^* and *Nab2 oe* brains are highly enriched for processes including RNA processing and neurodevelopment. **(A-C)** The enriched terms from FlyEnrichr database for Biological Process resulting from the subset of proteins increased in abundance that are **(A)** unique to *Nab2^ex3^*, **(B)** unique to *Nab2 oe*, and **(C)** shared between *Nab2^ex3^* and *Nab2 oe*. The bars shown correspond to the top ten c-scores (= ln(adj p-val) * z-score) in each dataset (adj. p-val<0.05). **(D)** A chord plot showing how proteins are represented in multiple GO Biological Process terms enriched from the subset of proteins increased in abundance in both *Nab2^ex3^* and *Nab2 oe* relative to control. The selected terms are shown on the right of the plot and are color coded according to the legend, with the chords extending to the left of the plot showing which proteins are represented in each term. The log_2_ fold-change in *Nab2^ex3^* is represented by color change (white to **red**) next to each protein annotation.

A similar analysis of shared and exclusive *decreased* proteins (**Figure 6A-D**) reveals that decreases unique to *Nab2^ex3^* are enriched in the processes of neuroblast proliferation, taste perception, and brain morphogenesis (**Figure 6A**), while *Nab2 oe* unique decreases are enriched in the processes of post-synapse assembly, synaptic vesicle recycling, and sodium ion transport (**Figure 6B**). The shared *decreases* between *Nab2^ex3^* and *Nab2 oe* represent processes involved in neurodevelopment and brain function (**Figure 6C**). A chord plot of biological process GO terms relating to neurodevelopment, behavior, and brain function highlights proteins *decreased* in both datasets (**Figure 6D**). Among these are the microtubule associated protein Futsch, the neuronal Ig-family transmembrane protein Turtle, the axon guidance and PCP component Vang, and the Rho GEF Trio (**Figure 6D**). In aggregate, these data provide a comprehensive view of the role Nab2 plays in regulating abundance of a specific cohort of proteins in the developing pupal brain, some of which are likely to correspond to mRNAs that are bound and regulated by Nab2 in brain neurons.

**Figure 6.**
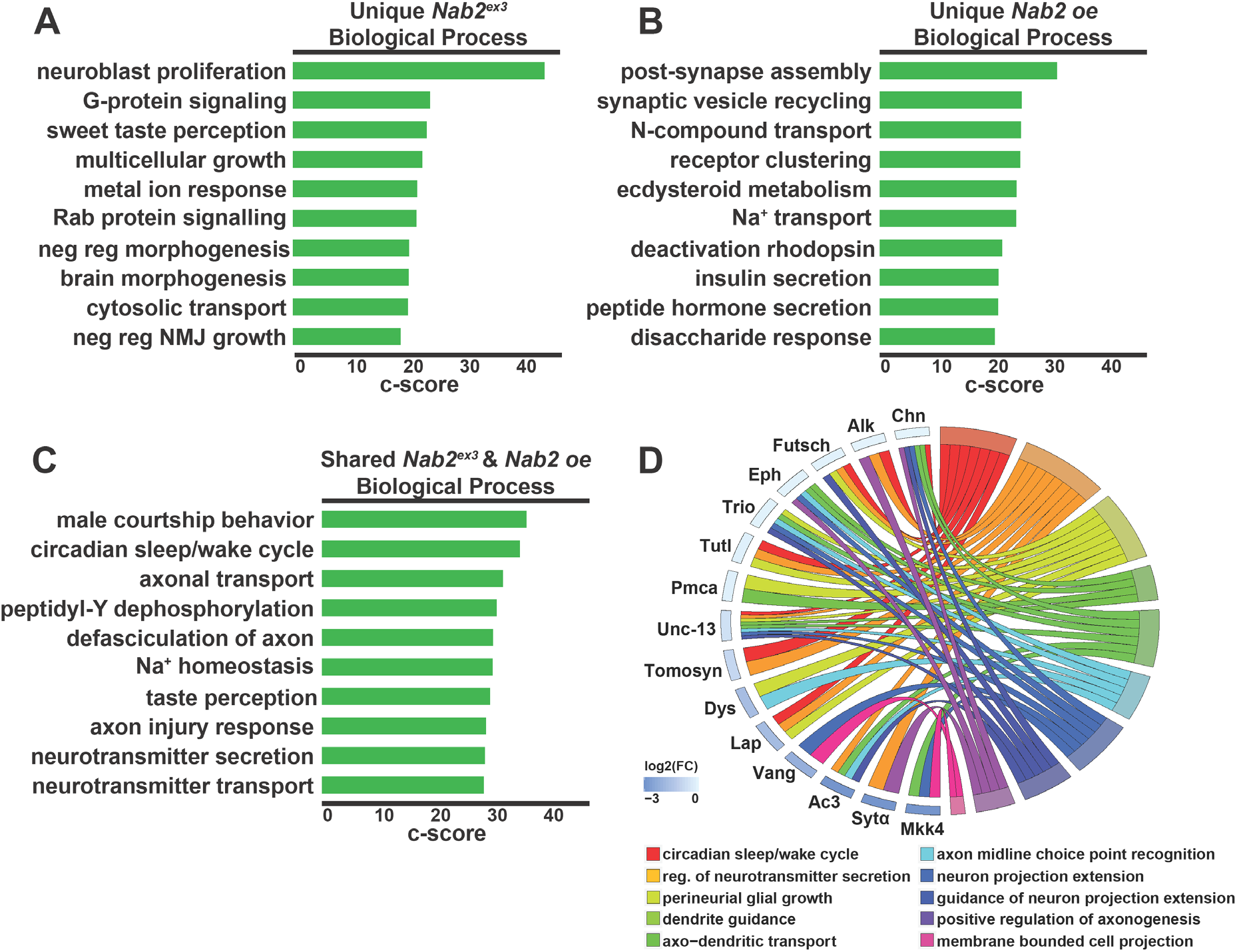
Proteins reduced in abundance in *Nab2^ex3^* and *Nab2 oe* are highly enriched for neurological roles. **(A-C)** The enriched terms from FlyEnrichr database for Biological Process resulting from the subset of proteins decreased in abundance that are **(A)** unique to *Nab2^ex3^*, **(B)** unique to *Nab2 oe*, and **(C)** shared between *Nab2^ex3^* and *Nab2 oe*. The bars shown correspond to the top ten c-scores (= ln(adj p-val) * z-score) in each dataset (adj. p-val<0.05). **(D)** A chord plot showing how proteins are represented in multiple GO Biological Process terms enriched from the subset of proteins decreased in abundance in both *Nab2^ex3^* and *Nab2 oe* relative to control. The selected terms are shown on the right of the plot and color coded according to the legend, with the chords extending to the left of the plot showing which proteins are represented in each term. The log_2_ fold-change in *Nab2^ex3^* is represented by color change (white to **blue**) next to each protein annotation.

## Discussion

Here, we characterize defects in the developing pupal brain and corresponding perturbations in the brain proteome caused by loss of the Nab2 RNA binding protein. Using carefully timed brain collections, we find that axon projection and development of MB α and β-lobes structure are more severely perturbed in pupal brains than in adult brains (α-lobe: 85% defective in pupal versus 50-60% in adult; β-lobe: 85% defective in pupal versus 70% in adult), and that coincident with these defects in axonal trajectories, we detect clear changes in a small fraction (~8%) of the brain proteome. This restricted effect on a subset of brain proteins is consistent with our recent finding that Nab2 loss has specific effects on the brain transcriptome (27), and supports the hypothesis that Nab2 regulates expression of a subset of neuronal mRNAs and proteins that are involved in various neurodevelopmental processes, including axon growth and guidance in the MBs.

Bioinformatic analysis of differentially expressed proteins in *Nab2^ex3^* mutant brains relative to control samples indicate that Nab2-regulated proteins are enriched in functional classes corresponding to axonal development, but also suggest a potential role in dendrites. The former link to axonogenesis matches the observed MB α and β-lobe defects, but the latter link to dendritic proteins is more novel and may be conserved. The murine Nab2 homolog, Zc3h14, localizes to dendritic shafts and spines and controls spine morphology in cultured neurons (21, 38). Nab2-regulated proteins identified in the current study that have predicted dendritic roles include the planar cell polarity factor Vang, the adhesion protein Cortactin, the netrin receptor Frazzled, the neuronal Ig-family transmembrane protein Turtle, the Fragile-X mental retardation homolog Fmr1, the Rho GEF Trio, the RNA binding protein Alan Shepherd/RBMS3, and the microtubule associated protein Futsch (MAP1β). Significantly, a proteomic dataset generated from hippocampi of P0 *Zc3h14* knockout mice (21) also shows enrichment for the Vang homolog Vangl2, in addition to five other neurodevelopmental proteins that are also detected here as differentially expressed in *Nab2^ex3^* pupal brains: the oxioreductase Wwox, the PDZ-domain protein X11Lβ/Apba1, the DnaJ protein CG6693/Dnajc9, the ARF-GEF factor Sec71/Psd3, and the endosomal protein Asrij/Ociad1 (**Table 1**). This evidence of conserved protein targets suggests that Nab2/ZC3H14 proteins may have a shared role in regulating key RNAs involved in neuronal development and signaling. Of note, fly Nab2 physically and functionally interacts with the *Drosophila* Fragile-X mental retardation protein (Fmrp1) (20), which has a key role in post-synaptic, activity-dependent local mRNA translation and is required for normal dendritic morphology (13).

Our comparison of the effects of Nab2 dosage reveals that almost one-third of proteomic changes (29%) that occur in Nab2-deficient pupal brains are shared in brains with neuronal overexpression of Nab2. Of 195 proteins that change in abundance in the *Nab2^ex3^* and *Nab2 oe* datasets, only 11 of these are inverse changes (i.e. ‘up’ in one genotype and ‘down’ in the other) while the other 184 proteins change in the same direction between these two genotypes. A simplistic model would predict that loss and gain of Nab2 would have the opposite effect on targets, but these data suggest that excess Nab2 can generate a dominant-negative effect on some candidate target RNAs, perhaps by sequestering Nab2-interacting proteins or blocking access of other RBPs to sites on RNAs. The remaining 11 proteins that show inverse changes in the *Nab2^ex3^* and *Nab2 oe* datasets could represent a subset of targets that respond in a linear fashion to Nab2 dose. One possibility is that the mRNAs encoding these proteins represent direct targets of the Nab2 RNA binding protein. Our data also detect 152 significantly changed proteins in *Nab2^ex3^* brains that are rescued back to normal levels in *Nab2 oe* brains, which parallels the morphological rescue of *Nab2^ex3^* by *Nab2 oe* documented in prior studies (15, 17, 29). Among the proteins in this group is Tumbleweed (Tum), which is homologous to human RacGap1 and required for normal MB development (39). This putative link from Nab2 to Tum-based control of MB patterning warrants further study.

Evidence of interactions between *Nab2* and elements of the microRNA (miRNA) machinery (e.g. *argonaute*) and ncRNA processing factors (e.g. *Rm62*) detected in our prior work (17, 20) are also supported by this proteomic analyses. Seven gene ontology (GO) terms relating to miRNA/ncRNA are enriched in the *Nab2^ex3^* dataset including *pre-miRNA processing, production of small RNA involved in gene silencing by RNA* and *ncRNA 3’-end processing*. As miRNAs and ncRNAs can regulate gene expression (40), some observed effects of *Nab2* alleles on the brain proteome could be indirect, rather than changes to direct (i.e. bound) Nab2 target RNAs. Our prior work indicates that Nab2 physically associates with the *Drosophila* homolog of the Fragile-X mental retardation protein, Fmr1, and coregulates some mRNAs (20). In the adult brain, depletion of Nab2 or Fmr1 each derepresses a *CamKII* translation reporter but has no effect on expression of a second Fmr1 target, *futsch* (20). In the current study of pupal brains, Futsch protein is decreased in *Nab2^ex3^* brains (log_2_(foldΔ) −0.38) while CamKII protein levels are not significantly changed. These stagespecific effects on the brain proteome raise the possibility that Nab2 interactions are not only target-specific (e.g. as in the case of alternative splicing) (27) but can also vary across developmental stages.

As noted above, the planar cell polarity (PCP) component Vang and its murine homolog Vang like-2 (Vangl2) are among a small group of homologous protein pairs that are differentially expressed in *Drosophila Nab2^ex3^* pupal brains and in P0 hippocampi dissected from *Zc3h14* knockout mice (21) (**Table 1**). This finding is particularly significant given the strong genetic interactions detected between an eye-specific *Nab2* overexpression system (*GMR-Nab2*) and multiple PCP alleles, including an allele of *vang* (41). The PCP pathway plays a conserved role in regulating axon projection and guidance in multiple higher eukaryotic species (42–46), including in the *Drosophila* MBs (47–51). Thus, the change in levels of Vang, a core PCP component (52–56), in *Nab2^ex3^* brains could provide an additional, direct link from Nab2 to a pathway that guides neurodevelopment including the MB α and β-lobes. In sum, the current study has revealed that *Nab2* loss perturbs the levels of a set of proteins that are enriched for neurodevelopmental factors that are, in turn, likely candidates to drive phenotypes in *Nab2* null neuronal tissue.

## Experimental Procedures

### *Drosophila* genetics

All crosses were maintained in humidified incubators at 25°C with 12hr light-dark cycles unless otherwise noted. The *Nab2^ex3^* loss of function mutant has been described previously (Pak et al., 2011). Alleles and transgenes: *Nab2^EP3716^* (Bloomington (BL) #17159) and *P{GawB}elavC^155^* (BL #458), and *w^1118^* (*‘control’*; BL #3605).

### Brain imaging, statistical analysis, and visualization

Brain dissections were performed as previously described (17). Briefly, 48-72 hours after puparium formation (APF) brains were dissected in PBS (1xPBS) at 4°C, fixed in 4% paraformaldehyde at RT, washed 3x in PBS, and permeabilized in 0.3% PBS-T (1xPBS, 0.3% TritonX-100). Following blocking for 1hr (0.1% PBS-T, 5% normal goat serum), brains were stained o/n in block+primary antibodies. After 5x washes in PBS-T (1xPBS, 0.3% TritonX-100), brains were incubated in block for 1hr, moved into block+secondary antibody for 3hrs, then washed 5x in PBS-T and mounted in Vectashield (Vector Labs). The anti-FasII monoclonal antibody 1D4 (Developmental Studies Hybridoma Bank) was used at 1:20 dilution. Whole brain anti-FasII images were captured on a Nikon AR1 HD25 confocal microscope using NIS-Elements C Imaging software v5.20.01, and maximum intensity projections were generated in ImageJ Fiji. Quantitation of MB phenotypes was performed as previously described (17).

### Global proteomics

#### Sample collection

Five biological replicates of control, *Nab2^ex3^*. and *Nab2 oe* for both female and male brains were collected at 23.25 – 25.5hr APF (5 pools per condition, 20 brains per pool), lysed in urea buffer (8M urea, 100mM NaHPO4, pH 8.5) with HALT protease and phosphatase inhibitor (Pierce) and processed at the Emory Proteomics Core.

#### LC-MS/MS

Data acquisition by LC-MS/MS was adapted from a previously published procedure (57). Derived peptides were resuspended in 20μL of loading buffer (0.1% trifluoroacetic acid, TFA). Peptide mixtures (2μL) were separated on a self-packed C18 (1.9μm, Dr. Maisch, Germany) fused silica column (25 cm x 75 μM internal diameter (ID); New Objective, Woburn, MA) and were monitored on an Orbitrap Fusion Tribrid Mass Spectrometer (ThermoFisher Scientific). Elution was performed over a 130-minute gradient at 250nL/min with buffer B ranging from 3% to 99% (buffer A: 0.1% formic acid in water, buffer B: 0.1% formic acid in acetonitrile). The mass spectrometer duty cycle was programmed to collect at top speed with 3s cycles. The full MS scans (300–1500 m/z range, 50ms maximum injection time) were collected at a nominal resolution of 120,000 at 200 m/z and AGC target of 200,000 ion counts in profile mode. Subsequently, the most intense ions above an intensity threshold of 5,000 were selected for higher-energy collision dissociation (HCD) (0.7 m/z isolation window with no offset, 30% collision energy, 10,000 AGC target, and 35ms maximum injection time) and the MS/MS spectra were acquired in the ion trap. Dynamic exclusion was set to exclude previously sequenced precursor ions for 30s within a 10ppm window. Precursor ions with charge states 2–7 were included.

#### MaxQuant

Label-free quantification analysis was adapted from a previously published procedure (58). Spectra were searched using the search engine Andromeda and integrated into MaxQuant against the *Drosophila melanogaster* Uniprot database (43,836 target sequences). Methionine oxidation (+15.9949 Da), asparagine and glutamine deamidation (+0.9840 Da), and protein N-terminal acetylation (+42.0106 Da) were variable modifications (up to 5 allowed per peptide); cysteine was assigned as a fixed carbamidomethyl modification (+57.0215 Da). Only fully tryptic peptides were considered with up to 2 missed cleavages in the database search. A precursor mass tolerance of ±20 ppm was applied prior to mass accuracy calibration and ±4.5 ppm after internal MaxQuant calibration. Other search settings included a maximum peptide mass of 6,000Da, a minimum peptide length of 6 residues, 0.6 Da tolerance for ion trap MS/MS scans. Co-fragmented peptide search was enabled to deconvolute multiplex spectra. The false discovery rate (FDR) for peptide spectral matches, proteins, and site decoy fraction were all set to 1%. Quantification settings were as follows: re-quantify with a second peak finding attempt after protein identification has completed; match MS1 peaks between runs; a 0.7 min retention time match window was used after an alignment function was found with a 20 minute RT search space. Quantitation of proteins was performed using summed peptide intensities given by MaxQuant. The quantitation method only considered razor plus unique peptides for protein level quantitation.

### Statistical analysis and data visualization

Statistical analyses were performed in either RStudio (Vienna, Austria) or GraphPad Prism 8 (Sand Diego, CA). Statistical analyses for MB phenotypes and plotting were performed using GraphPad. Significance determined using student’s t-test. Graphs reported either quartile ranks or error bars representing standard deviation. Significance scores indicated on graphs are * = p≤0.05, ** = p≤0.01, and *** = p≤0.001. Statistical analyses for proteomics data were performed using RStudio (59), custom in-house scripts, and the following packages: ggpubr (60), cluster (61), and GOplot (62). Gene ontology analyses were performed using FlyEnrichr (FlyEnrichr: amp.pharm.mssm.edu/FlyEnrichr/) (63–65).

### Data availability

The mass spectrometry proteomics data have been deposited to the ProteomeXchange Consortium via the PRIDE (66) partner repository with the dataset identifier PXD022984. All remaining data are contained within the article.

## Acknowledgements

We thank Dan Cox, GA State Neuroscience Institute, for reagents and discussion, and members of the Moberg and Corbett laboratories for helpful discussions. We thank the Emory Proteomics Core for their support and guidance.

## Funding and additional information

Research reported in this publication was also supported in part by the Emory University Integrated Cellular Imaging Microscopy Core of the Emory Neuroscience NINDS Core Facilities grant, 5P30NS055077. Financial support as follows: 5F31NS110312-02, 5F31HD088043-03, and 5R01MH107305-05.

## Conflict of Interest

The authors declare no competing financial interests.

## Notes

### Competing Interest Statement

The authors have declared no competing interest.

